# Synthetic analysis of natural variants yields insights into the evolution and function of auxin signaling F-box proteins in *Arabidopsis thaliana*

**DOI:** 10.1101/115667

**Authors:** R. Clay Wright, Mollye L. Zahler, Stacey R. Gerben, Jennifer L. Nemhauser

**Author notes:** Corresponding Author: Jennifer L. Nemhauser, HHMI Scholar, Department of Biology, University of Washington, Box 351800, Seattle, Washington 98195-1800 USA.

## Abstract

The evolution of complex body plans in land plants has been paralleled by gene duplication and divergence within nuclear auxin-signaling networks. A deep mechanistic understanding of auxin signaling proteins therefore may allow rational engineering of novel plant architectures. Towards that end, we analyzed natural variation in the auxin receptor F-box family of wild accessions of the reference plant *Arabidopsis thaliana* and used this information to populate a structure/function map. We employed a synthetic assay to identify natural hypermorphic F-box variants, and then assayed auxin-associated phenotypes in accessions expressing these variants. To more directly measure the impact of the strongest variant in our synthetic assay on auxin sensitivity, we generated transgenic plants expressing this allele. Together, our findings link evolved sequence variation to altered molecular performance and auxin sensitivity. This approach demonstrates the potential for combining synthetic biology approaches with quantitative phenotypes to harness the wealth of available sequence information and guide future engineering efforts of diverse signaling pathways.

## INTRODUCTION

Auxin controls many aspects of plant development and environmental adaptation. Natural and synthetic auxins have been used to control plant growth in fields, greenhouses and laboratories for nearly a century. In recent years, the gene families of biosynthetic and metabolic enzymes, transporters and perception machinery that determine the spatial, temporal and developmental specificity of auxin signals have been identified (Enders and Strader 2015). Recent work has just begun to determine how functionally robust the auxin signaling machinery is to mutation (Yu *et al.* 2013, 2015; Dezfulian *et al.* 2016), and to measure the propensity for mutations to produce novel plant phenotypes that result in evolutionary innovation (Delker *et al.* 2010; Rosas *et al.* 2013). As auxin effects are so wide-ranging, it is not surprising to find that significant variation exists in auxin sensitivity and auxin-induced transcription across *A. thaliana* accessions (Delker *et al.* 2010), perhaps contributing to morphological diversity. As such mapping evolutionary trajectories in auxin signaling could facilitate the engineering of numerous plant traits, such as root architecture, shoot branching or leaf venation—all traits associated with crop yield (Mathan *et al.* 2016).

Auxin is perceived by a coreceptor complex consisting of an F-box protein (TRANSPORT INHIBITOR RESPONSE1/AUXIN SIGNALING F-BOXES, TIR1/AFB; hereafter referred to as AFBs), an auxin molecule and a member of a transcriptional coreceptor/corepressor family (AUXIN/INDOLE-3-ACETIC ACID PROTEINS, Aux/IAAs). The F-box domain of the AFB associates with a Skp/Cullin/F-box (SCF) ubiquitin ligase complex that facilitates ubiquitination of the Aux/IAA proteins, targeting them for degradation (Lavy and Estelle 2016). In low auxin conditions, Aux/IAA proteins interact with and repress a family of transcription factors, the Auxin Response Factors (ARFs) (Guilfoyle and Hagen 2007). Auxin response genes are turned on when local auxin accumulation triggers degradation of Aux/IAAs thereby relieving the repression on ARFs.

*A. thaliana* has six *AFB* genes, *TIR1* and *AFB1*-*AFB5* (Dharmasiri *et al.* 2005a). The N-terminal F-box domain is modular and functionally conserved in TIR1 and AFB2, both of which form functional E3 ubiquitin ligase complexes with components in yeast and animals (Nishimura *et al.* 2009; Zhang *et al.* 2015). The C-terminal domain of the AFBs is a leucine-rich repeat (LRR). LRR domains offer a highly evolvable scaffold for binding small molecules and proteins and perform diverse functions across all domains of life (Bella *et al.* 2008). The AFB LRR domain allows auxin sensing by interacting with both auxin and the Aux/IAA transcriptional repressor/co-receptor proteins (Dharmasiri *et al.* 2005a; Tan *et al.* 2007; Calderón Villalobos *et al.* 2012). The identity of the subunits and their affinity for one another governs the rate of Aux/IAA degradation which, in turn, governs transcriptional dynamics, cell fate and morphological change (Dreher *et al.* 2006; Pierre-Jerome *et al.* 2014; Guseman *et al.* 2015; Galli *et al.* 2015).

Here, we paired an examination of the natural coding sequence variation in the AFB family with quantification of functional variation. We used a synthetic auxin-induced degradation assay in yeast to assess the function of natural variants in isolation from the rest of the auxin response network. Variants with altered function were then evaluated in their native context by quantifying auxin-associated root growth inhibition in accessions containing these polymorphisms. Finally, we directly measured the contribution to auxin sensitivity of the most hypermorphic *TIR1* allele by generating transgenic plants expressing this variant under a constitutive promoter. Through this work, we have generated a higher resolution structure/function map of the AFB family and highlighted the challenge of identifying functional divergence in highly buffered signaling pathway components using intact plants.

## MATERIALS AND METHODS

### Materials, media composition and general growth conditions

PCRs were performed with Phusion (cloning reactions; NEB, Ipswich, MA), GoTaq (diagnostics; Promega, Madison, WI) or GemTaq (genotyping; MGQuest, Lynnwood, WA) with primers from IDT (Coralville, Iowa). Media were standard formulations as described in (Pierre-Jerome *et al.* 2017). Plants were grown on 0.5x LS media (Caisson Laboratories, Smithfield, UT) containing 0.5% sucrose and 0.7% phytoagar (plantmedia, Dublin, OH). Seeds were obtained from the Arabidopsis Biological Resource Center (Columbus, OH).

### Analysis of sequence variation

A reference dataset of the genome locations of the TIR1/AFB family and COI1 was assembled from the TAIR10 database on 28 July 2015. Transcript and coding sequences were identified using the ENSEMBL biomart version of TAIR10. The 1001 genomes Salk dataset (28 June 2010) was obtained from http://1001genomes.org/. Single nucleotide polymorphisms (SNPs) and one base pair deletions with a quality (PHRED) score of 25 and above (i.e. “quality_variant_filtered” files) were used for the following analysis using a custom R scripts unless otherwise specified. SNPs located in genes of interest were isolated and mapped to their respective gene structures using the VariantAnnotation package (Obenchain *et al.* 2014). Coding variants were identified and assembled for each gene and each accession. Nucleotide diversity, Watterson’s theta and Tajima’s D were calculated using the PopGenome package (Pfeifer *et al.* 2014).

In identifying polymorphisms, TIR1/AFB genes were split into F-box and LRR domains, with the F-box defined as the N-terminus of the protein to I50 of TIR1 and the corresponding residues of the other genes according to the alignment generated by Tan et al. (Tan *et al.* 2007). The N-terminal extension of AFB4 and 5 were excluded.

For functional analysis, nonsynonymous polymorphisms in *TIR1* and *AFB2* were isolated. As domain swap experiments revealed that the F-box regions of TIR1 and AFB2 confer highly similar or identical function in yeast (S4 Fig), we focused our analysis on variants within the LRR domain. Highly represented and potentially functionally divergent nonsynonymous polymorphisms were then identified by creating a dN/dS matrix of all-by-all pairs of accessions for each gene using the kaks function within the seqinr R-package (Charif and Lobry 2007), which implements the method of Nei and Gojobori (Nei and Gojobori 1986). Incalculable and infinite values were excluded from these matrices prior to extraction of outlier pairs and associated nonsynonymous polymorphisms. This set of TIR1 and AFB2 polymorphisms was then cloned into yeast expression vectors and functionally characterized as described below. The remaining polymorphisms were subsequently cloned and characterized (Fig S6 and S7). Annotated code and supplemental data are in S11 Appendix.

### Strain construction

Plasmids were designed using j5 (Hillson *et al.* 2012) and constructed by aquarium (www.aquarium.bio). TIR1 and AFB2 were separately inserted into pGP8G (Havens *et al.* 2012) downstream of a GPD promoter and followed by 3X-FLAG-6X-HIS tandem affinity purification tag, via Golden Gate cloning (Engler *et al.* 2009). Mutations were introduced into the parent vectors via two-fragment Gibson assembly (Gibson *et al.* 2009). The coding sequence of the gene of interest was confirmed by sequencing (Genewiz, South Plainfield, NJ).

Plasmids were digested with *PmeI* before Lithium PEG (37) transformation into W303-1A ADE2+ yeast (MATa, leu2-3,112 trp1-1 can1-100 ura3-1 his3-11,15 ybp1-1). Correct integration of transformed colonies was confirmed by diagnostic PCR across the 3’ boundary of homologous recombination, relative to the gene of interest. Similarly, pGP4GY-IAA1 and -IAA17 (Havens *et al.* 2012) were transformed into W814-29B yeast (MATα ade2-1 trp1-1 can1-100 ura3-1 leu2-3,112 his3-11,15). Confirmed transformants were struck to isolation on YPAD plates. *AFB* strains were individually mated with each *Aux/IAA* strain using standard methods (Pierre-Jerome *et al.* 2016).

### Auxin-induced degradation assays in yeast

Assays were essentially as described in (Pierre-Jerome *et al.* 2017) using a BD special order cytometer with a 514 nm laser exciting fluorescence that is cutoff at 525 nm prior to PMT collection (Becton-Dickinson, Franklin Lakes, NJ). Events were annotated, subset to singlet yeast, and normalized to initial levels of fluorescence using the flowTime R package (http://www.github.com/wrightrc/flowTime). Full dataset is available via FlowRepository (http://tinyurl.com/j268y5e). Additional detail in S11 Appendix.

### Western blot analyses

Yeast cultures that had been incubated overnight in SC media were diluted to OD_600_ = 0.6 and incubated until cultures reached OD_600_ ~ 1. Cells were harvested by centrifugation from four milliliters of culture. Cells were then lysed by vortexing for five minutes at 4°C in the presence of 100 µL of 0.5 mm diameter acid washed glass beads and 200 µL SUMEB buffer (1% SDS, 8 M urea, 10 mM MOPS pH 6.8, 10 mM EDTA, 0.01% bromophenol blue) per 1 OD unit of original culture. Lysates were then incubated at 65°C for ten minutes and cleared by centrifugation prior electrophoresis and blotting (Sambrook and Russell 2001). Mouse anti-FLAG M2 monoclonal primary antibodies (Sigma-Aldrich, St. Louis, MO) were used at a 1:1000 dilution per the manufacturer’s directions.

### Root growth inhibition assays

After sterile seeds were stratified on plates oriented vertically at 4°C in the dark for 3 days (or 1 week for wild accessions), they were transferred to long day conditions at 20°C for 4 days. Ten plants each of 4 different genotypes were then transferred in two rows to plates containing either DMSO carrier or 2,4-dichlorophenoxyactic acid (2,4-D) with root tips aligned to a reference mark for each row. Plants were scanned after an additional 3 days of incubation. Plates were regularly rotated during incubation to avoid position effects. Root growth was measured using ImageJ (Rasband 1997) and an Intuos Pro drawing pad (Wacom, Portland, Oregon). Additional detail can be found in S11 Appendix. The experiment in Fig 3A-C was repeated on two different days. The experiments in Fig 3D were repeated on three different days for T2 lines and five different days to T3 lines.

### Construction and analysis of transgenic plants

Genes of interest were inserted via Golden Gate cloning (Engler *et al.* 2009) into pGreenII (Hellens *et al.* 2000) with a pUBQ10 promoter (Grefen *et al.* 2010) and 3X-FLAG-6X-HIS tandem affinity purification tag. Plasmids were transformed into *Agrobacterium tumefactions* GV3101 containing pSOUP (Hellens *et al.* 2000) via electroporation, and transformants were selected on plates with 50 µg/mL gentamycin and 25 µg/mL kanamycin. Plants were transformed by floral dip (Zhang *et al.* 2006), and transformants were selected on plates with 30 µg/mL hygromycin at four days post germination after an initial light exposure for seven hours. Root growth inhibition phenotypes were quantified in T2 generation of three independent transformants as described above. Each plant was genotyped for the presence of the hygromycin resistance gene after the growth assay, using the forward primer (GATGTTGGCGACCTCGTATT) and the reverse primer (GTGCTTGACATTGGGGAGTT). Expression levels in T3 lines were measured by quantitative PCR. RNA was isolated from the tissue from young leaves of T3 plants using illustra RNAspin Mini RNA Isolation Kit (GE Healthcare, Little Chalfont, United Kingdom) and reverse transcribed using iScript cDNA synthesis kit (Bio-rad, Hercules, California). Quantitative PCR was performed using iQ SYBR Green supermix (Bio-rad, Hercules, California) and primers for *TIR1* (f-CACGGAACAAGAAGACATCCAAAGG, r-TGAGGAAACTAGAGATAAGGGACTGC) or *PP2A* (f-AACGTGGCCAAAATGATGC, r-AACCGCTTGGTCGACTATCG) in a CFX96 Touch Real-Time PCR detection system (Bio-rad, Hercules, California).

Plasmids, strains and sequence files are available upon request or via Addgene. All code used to perform analysis and visualization is provided in S11 Appendix. All data including raw images are available upon request.

## RESULTS

We identified polymorphisms across the entire AFB gene family in the 170 *A. thaliana* accessions of the SALK subset of the 1001 Genomes Project (Schmitz *et al.* 2013). The AFB gene family is highly conserved relative to other auxin signaling gene families (Delker *et al.* 2010). We found 1,631 total polymorphisms within coding regions and 175 segregating sites across the whole family (S1 Table and S2 Fig). *AFB3* had the highest ratio of per site diversity at nonsynonymous sites relative to synonymous sites. *AFB4*, critical for response to the synthetic auxin picloram (Prigge *et al.* 2016), had the highest nonsynonymous diversity (more than 10X that of *TIR1*) and the only two nonsense polymorphisms identified in this dataset. In contrast, *AFB1*, which is largely incapable of forming a functional SCF complex (Yu *et al.* 2015), has similar ratio of nonsynonymous to synonymous diversity as *TIR1*. Many of the accessions contained nonsynonymous polymorphisms in multiple members of the AFB family (S5 Table). These additional polymorphisms occurred more frequently in TIR1/AFB1 and AFB2/AFB3 sister pairs than expected (permutation analysis, p<0.05, S11 Appendix section “Assessing covariation…”).

None of the identified accessions have nonsynonymous polymorphisms in both *TIR1* and *AFB2* (S5 Table), an unlikely pattern to occur by chance (permutation analysis, p<0.05, S11 Appendix section “Assessing covariation…”). This may reflect the fact that *AFB2* and *TIR1* are the major auxin receptors and serve partially redundant functions, a conclusion supported by genetic analysis (Dharmasiri *et al.* 2005a; Parry *et al.* 2009). The majority of the nonsynonymous polymorphisms in TIR1 and AFB2 occurred in positions of high diversity across the Col-0 AFB family, and most were located in surface residues of the LRR domain (Fig 1A). The majority of these polymorphisms spanned the exterior helices and loops of the fourth through eighth LRRs, which face the Cullin subunit (Fig 1B and 1C). This region was recently identified as being responsible for SCF^TIR1^ dimerization (Dezfulian *et al.* 2016) and is also proximal to the S-nitrosylation site (Terrile *et al.* 2012). A pair of polymorphisms exists on the surface spanning the final three LRRs and the C-terminal cap (Fig 1D). This region may interact with the KR motif known to strongly affect auxin-induced degradation rates (Dreher *et al.* 2006; Moss *et al.* 2015). A final pair of polymorphisms was found on the interior surface of the LRR domain horseshoe (Fig 1E).

**Fig 1.**
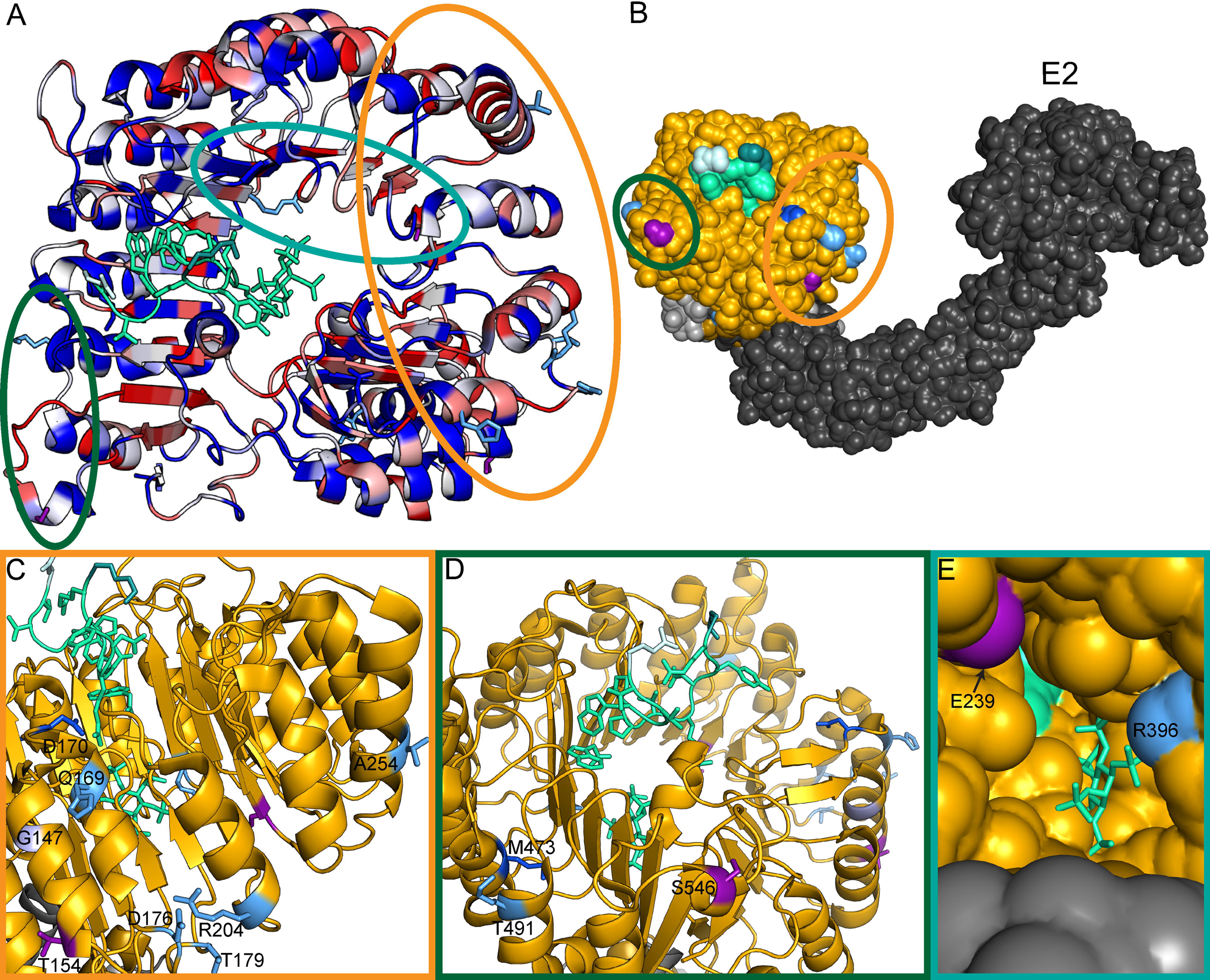
Clusters of natural variation in *TIR1* and *AFB2*. (A) Identified nonsynonymous polymorphisms tend to occur in residues of high diversity within the *Arabidopsis* AFB family. A top down view of the LRR domain of the TIR1 structure (PDB:2P1Q) is shown with the F-box domain in the bottom right and the LRR domain spiraling counterclockwise. The backbone of the TIR1 structure (Tan *et al.* 2007) was colored according to protein sequence diversity with conserved residues in blue and diverging residues in red. Diversity was calculated as Shannon Entropy using an alignment of the protein sequences of the *Arabidopsis* AFB family (TIR1, AFB1-5). All nonsynonymous polymorphisms are shown as sticks. AFB2 variants are in light blue and TIR1 variants are in purple. Previously identified TIR1 mutations are in dark blue (Ruegger *et al.* 1998; Yu *et al.* 2013). The IAA7 degron is shown as a light green ribbon with side-chains as sticks. The N-terminal residue of the IAA7 degron is in lighter green and the C-terminal residue is darker green. Circles around polymorphisms match the detailed views shown in panels C, D and E. (B) Polymorphisms face the Cullin subunit of the predicted SCF^TIR1^ structure. ASK1 (light grey) was aligned with SKP1 from the human SKP2-SKP1-Cul1-RBX1 structure (PDB: 1LDK, shown in dark grey), docking with TIR1 (gold). Putative E2 location is labeled. (C) The dimerization domain on the N-terminal side of the LRR horseshoe contains the majority of natural variation in *TIR1* and *AFB2*. The *tir1-1* allele (*tir1^G147D^*) is in light purple. (D) Two variants were located on the C-terminal side of the LRR close to the N-terminus of the degron. (E) Two additional variants were located inside the LRR horseshoe, near the inositolhexakisphosphate cofactor.

### Synthetic yeast assays reveal functional variation in TIR1 and AFB2

An auxin-induced degradation assay has been established in yeast using heterologous expression of either *TIR1* or *AFB2* (Havens *et al.* 2012). We used this synthetic assay to quantify the function of AFB natural variants in the absence of the potentially confounding effects of feedback from the auxin pathway itself or from modulation by other integrating pathways. Natural variants were engineered into the Col-0 reference sequence with co-occurring polymorphisms cloned individually and in combination. Each AFB was then constitutively co-expressed in yeast with fluorescently labeled Aux/IAA targets. Auxin-induced degradation was measured for two targets, IAA1 and IAA17, as these substrates show distinct patterns of behavior when assayed with Col-0 TIR1 and AFB2. TIR1^Col^ induces degradation of IAA1 and IAA17 at similar rates, while AFB2^Col^ causes IAA17 to degrade much faster than what is observed for IAA1 (Havens *et al.* 2012). We focused on polymorphisms in the LRR domain that were predicted to be functionally divergent (having any pairwise *d_N_*/*d_S_* value greater than one), but analysis of the few additional polymorphisms is shown in Figures S6 and S7.

Some natural variants increased function compared to the Col-0 reference, while others decreased or nearly abrogated function (referred to hereafter as hypermorphs, hypomorphs and amorphs, respectively) (Fig 2). Of the *TIR1* polymorphisms, T154S was hypermorphic and E239K-S546L was strongly hypomorphic (Fig 2A). E239K alone was nearly amorphic, and adding S546L only slightly restored activity. These polymorphic TIR1 variants are expressed at similar levels to TIR1^Col^ (S8 Fig). Among the *AFB2* polymorphisms, T491R was the only clear hypermorph identified (Fig 2B). D176E was slightly hypermorphic, whereas A254V was a moderate hypomorph. In combination, these two polymorphisms were largely additive, giving a response quite similar to AFB2^Col^. AFB2^Q169L^ was also a moderate hypomorph. Two *AFB2* alleles, R396C and R204K, were strong hypomorphs, and T179M was amorphic in our assays. Interestingly, the two most highly represented variants, TIR1^T154S^ (present in 5 accessions) and AFB2^R204K^ (6 accessions), show strong functional divergence from their respective wild-type proteins.

**Fig 2.**
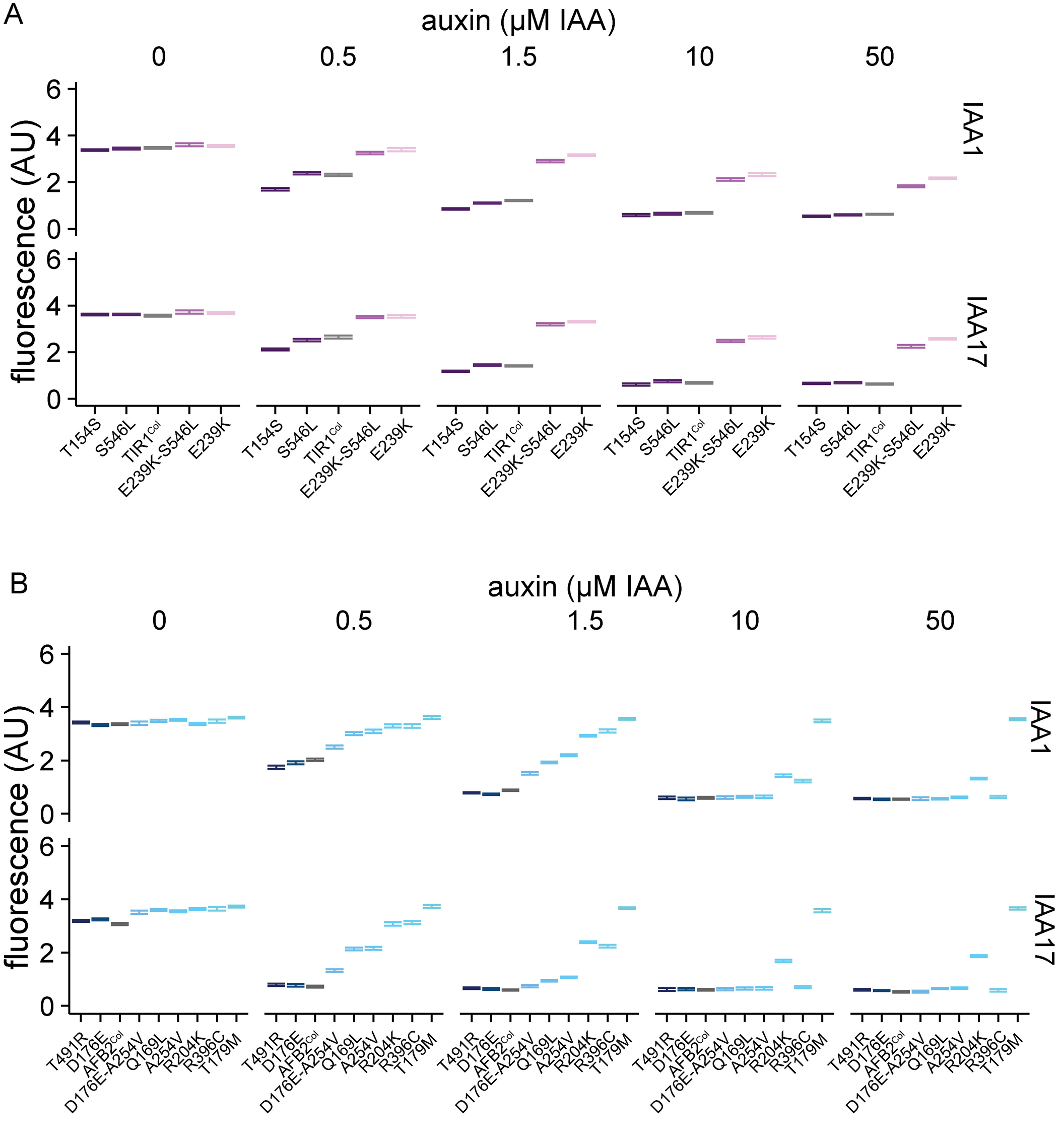
Synthetic assays reveal significant functional variation in naturally occurring *AFB* polymorphisms. Nonsynonymous polymorphisms in the LRR domains of *TIR1* (A) and *AFB2* (B) were synthesized and co-expressed in yeast with fluorescently labeled IAA1 or IAA17. Degradation was assessed using flow cytometry on cultures exposed to different concentrations of the auxin indole-3-acetic acid (IAA) for one hour. Error bars represent 95% confidence intervals around the median fluorescence calculated from three independent experiments. In many cases, intervals are small enough that they appear as a single line. The reference Col-0 variant is shown in grey.

### Accessions containing a hypermorphic *TIR1* allele are hypersensitive to auxin

We next assessed whether the functional variation observed in the synthetic assays was manifested as phenotypic differences in the respective accessions. As there are many polymorphisms between accessions and previous genome-wide association studies of auxin response have not identified the *AFB* family (Rosas *et al.* 2013; Meijón *et al.* 2014), we did not expect a strong correlation between genotype and phenotype from our analysis. To increase the sensitivity and precision of the test, we measured inhibition of primary root growth in the presence of exogenous auxin and fit a log-logistic dose response model to the data. Similar bioassays have been used extensively to identify and characterize mutants in the *AFB* gene family (Gray *et al.* 1999; Dharmasiri *et al.* 2005a; b; Parry *et al.* 2009). The effective dose of auxin required to elicit fifty percent of the maximum root growth inhibition (ED50) was the most effective parameter in our model for differentiating among genotypes. Two *tir1* mutants in the Col-0 background (a point mutation *tir1-1* and a T-DNA insertion *tir1-10*) were also included in our analysis. Both mutants had significantly higher ED50 values than Col-0, as expected (Fig 3A and C). A loss of function *afb2* allele did not significantly affect the root growth response in our assays, although *tir1-1 afb2-3* double mutants had a much larger ED50 than the *tir1-1* single mutant.

**Fig 3.**
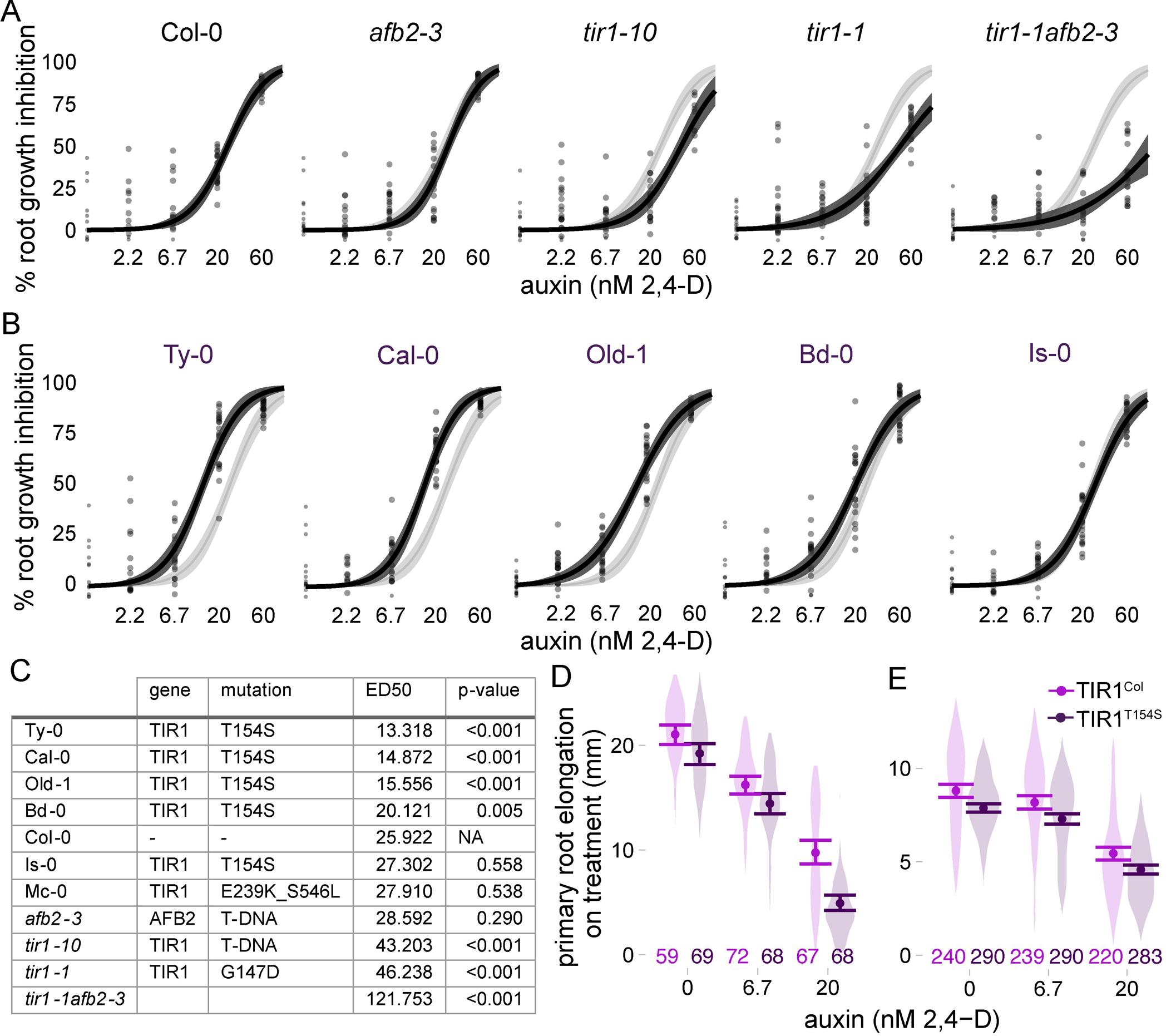
Auxin sensitivity varies only subtly in wild accessions. (A and B) The impact of auxin on root growth (normalized to mock treated controls) was measured in 8-day-old seedlings. Results from two biological replicates, each containing ten plants per treatment level, are shown. Each measurement is shown as a transparent grey point. Solid lines represent log-logistic dose response model fits with a lighter ribbon indicating 95% confidence intervals. The Col-0 curve is reproduced in light grey in each panel to facilitate comparisons. (A) Assays on the reference accession Col-0, and mutants in the Col-0 background, are shown (B) Auxin sensitivity of accessions containing the hypermorphic *TIR1^T154S^* allele. (C) Estimated ED50 values for selected accessions and controls. Parameters were compared ratiometrically to Col-0 and one-sample t-tests were used to estimate the likelihood that the ratio of parameters equals 1. P-values were corrected for multiple testing using the Benjamini-Hochberg method. (D and E) A natural polymorphism was sufficient to alter auxin sensitivity in plants. Mean root growth (large points) and 95% confidence intervals (error bars) are shown on top of a violin plot representing the distribution of all measurements. All transgenes were expressed under the pUBQ10 promoter. The number of plants measured for each condition is shown above the X-axis with individual measurements indicated by small points. (D) Three experimental replicates were performed with T2 plants from four independent lines of Col-0 expressing the reference allele and three independent lines expressing *TIR1^T154S^*. (E) Five experimental replicates were performed with T3 plants from five independent lines of Col-0 expressing the reference allele and six independent lines expressing *TIR1^T154S^*.

Four out of five accessions carrying *TIR1^T154S^* were hypersensitive to auxin, following the pattern predicted by the hypermorphic behavior of that variant in yeast (Fig 3B and C). In three of these accessions, ED50 values were quite close to one another and significantly lower than the ED50 measured for Col-0 or any other accession. However, Mc-0, which contains the strong hypomorph *TIR1^E239K-S546L^*, had auxin responses that were only subtly different Col-0. Consistent with the modest phenotype of *afb2-3* in our assays, auxin responses of most accessions with *AFB2* polymorphisms were essentially similar to those of Col-0 (S11 Appendix, pg. 38-40).

### A common *TIR1* allele confers auxin hypersensitivity to Col-0

The aberrant auxin responses in yeast and the majority of accessions led us to hypothesize that *TIR1^T154S^* is a natural gain-of-function allele with the capacity to impact organ-level auxin responses. To test this, we generated transgenic Col-0 lines expressing *TIR1^Col^* or *TIR1^T154S^* under a constitutive promoter. Most transgenic lines had relatively similar expression levels, although the Col-0 allele was expressed on average at modestly higher levels than the T154S variant (Fig S9). Expression level and root growth phenotypes were not strongly correlated—the lines with the lowest expression levels showed essentially similar auxin responses as lines with higher transgene expression. In the T3 generation, silencing of the endogenous *TIR1* and transgenes was observed in three lines. Two of these lines were from the family with highest transgene expression (Fig S9, TIR1-7).

Overall, plants expressing *TIR1^T154S^* had shorter roots than plants expressing *TIR1^Col^* even in the absence of auxin treatment, consistent with the expectation that the T154S polymorphism conferred increased sensitivity to endogenous auxin. In the T2 generation, auxin treatments in root inhibition assays confirmed that *TIR1^T154S^* had increased auxin sensitivity relative to *TIR1^Col^* (Fig 3D) (transgene:treatment effect F = 9.3, p = 0.0001, full statistical analysis shown in S11 Appendix). In the T3 generation, *TIR1^T154S^* expressing plants consistently had shorter roots than *TIR1^Col^* expressing plants (transgene effect F = 100.4, p < 10^−16^); however, the difference in sensitivity to exogenous auxin when compared with *TIR1^Col^* expressing plants was diminished. This may be because the auxin response is near saturation even in the *TIR1^Col^* expressing plants in this generation. The roots of homozygous T3 plants with either transgene were much shorter and had a significantly reduced auxin response when compared with the T2 generation (Fig 3D vs. E, note especially the difference in the y-axis).

### Dimerization domain variation affects dominance relations between TIR1 alleles

One of the unexpected findings in our analysis of auxin response across genotypes was a subtle but highly reproducible difference between the two induced alleles of *tir1* in the Col-0 background (Fig 3A, C). The point mutation *tir1-1* showed a consistently stronger loss of auxin sensitivity than the T-DNA insertion *tir1-10*, raising the possibility that *tir1-1* might be acting as a dominant negative or antimorph rather than as a simple loss-of function. In support of that interpretation, *tir1-1* mutants are semi-dominant (Ruegger *et al.* 1998), and the *tir1-1* allele (G147D) and several other mutations in nearby residues negatively affect SCF^TIR1^ dimerization and activity (Dezfulian *et al.* 2016).

We turned to the yeast synthetic system to further investigate this question. By transforming a single copy of each allele into haploid yeast strains of each mating type, we created all pairs of alleles via mating. We also created *tir1*^K159*^a mimic of the *tir1-10* T-DNA insertion allele. As expected, *tir1*^K159*^ was an amorph, behaving similarly to an empty expression cassette (S10 Fig). *TIR1* dosage had little effect on auxin response in these assays, as *TIR1/tir1-10* heterozygotes responded similarly to *TIR1* homozygotes (Fig 4A). In contrast, expression of *tir1-1* nearly completely abrogated *TIR1* activity (Fig 4B), providing strong evidence that *tir1-1* is a dominant negative allele. In addition to having a greatly reduced ability to induce Aux/IAA degradation, it is likely that *tir1-1* is also outcompeting TIR1 for SCF complex formation and substrate binding as *tir1-1* protein accumulated to much higher levels in yeast than *TIR1* (Fig 4C).

**Fig 4.**
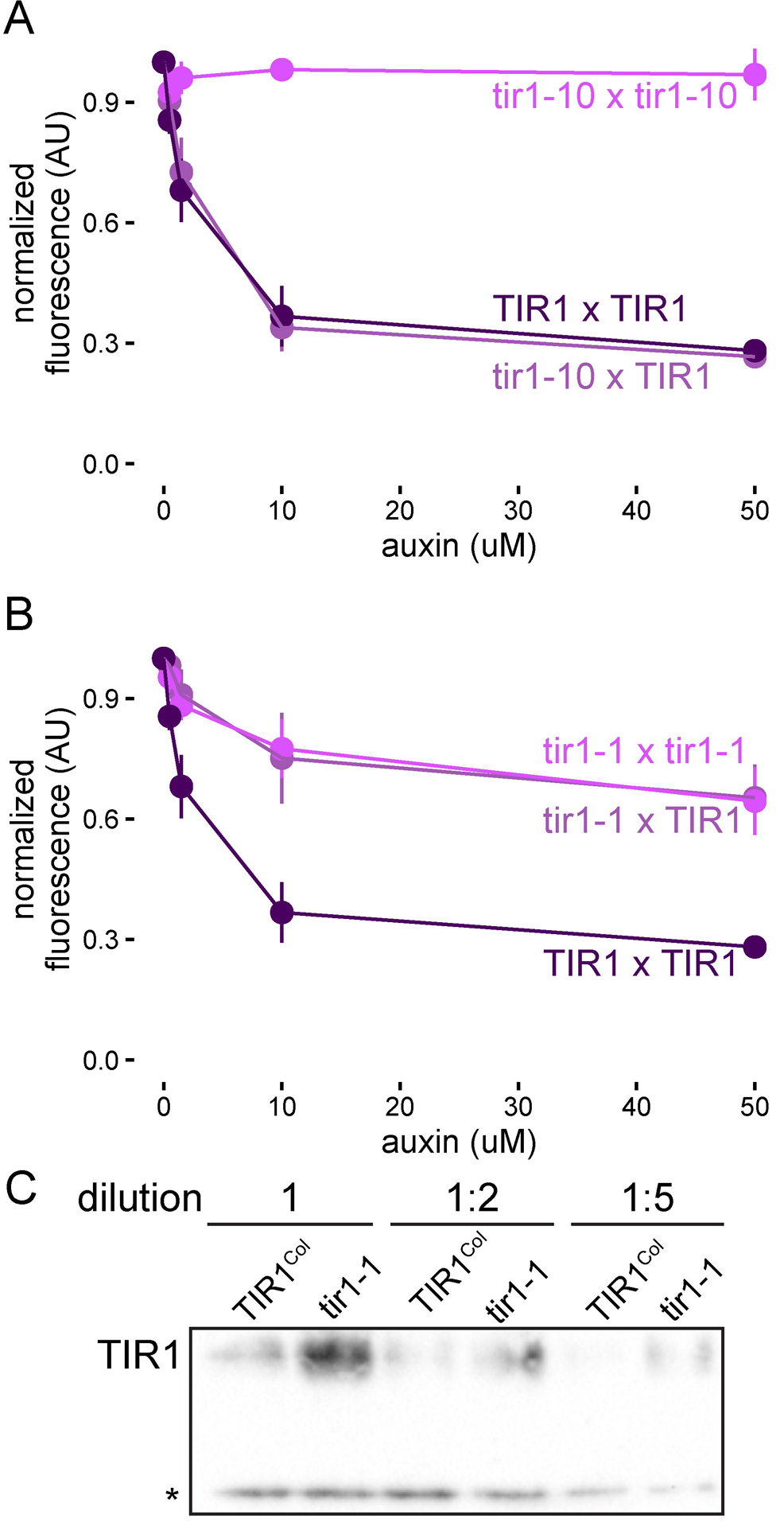
tir1-1 is a dominant negative allele. Yeast expressing YFP-IAA17 and pairwise combinations of (A) *TIR1* and *tir1-10* (*tir1^K159*^*) or (B) *TIR1* and *tir1-1* (*tir1^G147D^*) alleles were treated with various concentrations of auxin for one hour before YFP-IAA17 fluorescence was measured by flow cytometry. Mean fluorescence +/- SE was calculated from four experiments. Some error bars are within the points.

## DISCUSSION

The analysis of intraspecific variation in auxin sensitivity presented here critically extends previous work on the evolution of this pathway by focusing on protein level functional variation. Synthetic assays allowed for direct quantification of differences in the ability of TIR1 and AFB2 variants to facilitate ubiquitin-mediated degradation of their substrates. The creation of a structure/function map of natural variation revealed several areas of the F-box-LRR protein scaffold that can accommodate mutations, while modulating auxin sensitivity. For example, this analysis further underscored the importance of the AFB dimerization domain (Dezfulian *et al.* 2016) to regulate SCF activity.

The *AFB* family provides a test case for genome evolution after gene duplication, as there is evidence of both significant novelty and redundancy between family members (Dharmasiri *et al.* 2005a; Walsh *et al.* 2006; Parry *et al.* 2009; Hu *et al.* 2012). Analysis of intraspecific coding sequence polymorphisms in this study has identified a subset of the tolerated polymorphisms within the AFB family. This natural variation has also revealed potential differences in evolutionary rates across the gene family and redundancies within sister pairs. In the future, quantitative phenotyping and precision genetics will allow us to test related hypotheses and accurately partition the novel and redundant effects of individual AFB genes on developmental phenotypes. Accessions containing highly represented polymorphisms and having phenotypes not predicted by our synthetic functional analysis, should facilitate future examination of evolutionary robustness and plasticity in nuclear auxin signaling and downstream gene networks.

Our analysis demonstrates that functional diversification is occurring within the *Arabidopsis TIR1* lineage and clarified the role of induced variants that are commonly used for auxin studies. The integrated biochemical and phenotypic analysis of natural variants refined the map of functionally relevant residues in TIR1 and AFB2. Synthetic analysis of the chemically-induced *tir1-1* allele, which is in proximity to many of the natural polymorphisms found in *TIR1* and *AFB2*, has established *tir1-1* as a dominant negative allele and highlighted the potential importance of interactions among family members to the auxin response.

The auxin pathway in *Arabidopsis*, like many critical signaling pathways across eukaryotes, has high levels of redundancy at each node and numerous modes of feedback. Together these factors act as strong buffers masking functional changes in any one component. This effect likely explains the discrepancies between synthetic and plant phenotypes in this study. Similar factors may also contribute to the lack of *TIR1/AFB* genes identified in genome-wide association studies. Candidate gene approaches incorporating isolated functional assays as demonstrated here can complement genome-wide approaches by removing feedback and other compensatory effects. Future efforts that combine synthetic assays with higher throughput allelic replacement technologies in plants (Čermák *et al.* 2017) would substantially increase the ability to precisely compare the impact of a given variant in isolation and in a common plant context.

Extending this pipeline for structure/function and genotype/phenotype mapping to additional auxin signaling genes and developmental phenotypes will improve our understanding of how plant form is shaped by this small molecule. This information, along with the general evolvability of the LRR scaffold (Bella *et al.* 2008), make the AFBs potential candidates for engineering novel traits in crops (Sun *et al.* 2016).

## ACKNOWLEDGEMENTS

We thank Doug Fowler, Adam Leaché and Eric Klavins for guidance on methods, analysis and interpretation of our findings; members of the Nemhauser, Klavins and Imaizumi Labs for helpful discussions; and Brenda Martinez for technical assistance. This work was supported by the National Institute of Health (R01-GM107084), the National Science Foundation (MCB-1411949) and the Howard Hughes Medical Institute. R.C.W. received fellowship support from the National Science Foundation (DBI-1402222).

## SUPPORTING INFORMATION CAPTIONS

**S1 Table. Sequence variation in the AFB gene family.**

**S2 Fig. Polymorphisms in the *AFB* genes of the 170 analyzed accessions.** Using a sliding 5-codon window, synonymous (blue dotted) and nonsynonymous (red solid) diversity per site was calculated across each AFB gene for all 170 accessions. A vertical black dotted line separates the F-box and LRR domain of each gene and also identifies the target site of miR393. Only those genes strongly affected by miR393 are labeled. Nonsynonymous polymorphisms functionally characterized in this study are indicated.

**S3 Fig. Map of *TIR1* and *AFB2* polymorphic accessions.** The 170 resequenced accessions analyzed in this study were mapped to the latitude and longitude of their reported collection sites. Each accession is shown as a transparent grey diamond, such that locations with many accessions appear darker grey. Accessions containing functionally characterized nonsynonymous polymorphisms in *TIR1* or *AFB2* are highlighted in pink/purple and cyan/navy respectively, with darker tones representing hypermorphs and lighter tones representing hypomorphs. (A) Many *TIR1 or AFB2* polymorphic accessions were collected across Europe. (B) A cluster of *AFB2* polymorphic accessions was collected around the Great Lakes region of the United States of America.

**S4 Fig. Functional divergence between TIR1 and AFB2 results from sequence divergence in the LRR domains.** (A) Schematics of domain swaps between TIR1 and AFB2. Amino acid ranges for each domain are labeled. Line type legend for panel B is to the left of the schematics. (B) Auxin-induced degradation rates are encoded by AFB LRR and not F-box domains. Degradation rates of a fluorescently-labeled IAA17 substrate were measured in yeast expressing wild-type or domain-swapped variants of TIR1 and AFB2 and treated with auxin. Data from two independent experiments is shown. Mock-treated controls are shown in grey. Wild-type TIR1 and TIR1 with an AFB2 F-box domain are indicated in purple, shown as solid and dotted lines respectively. Wild-type AFB2 and AFB2 with a TIR1 F-box domain are indicated in blue, shown as solid and dotted lines respectively.

**S5 Table. Accessions containing nonsynonymous variants in *TIR1* or *AFB2.***

**S6 Fig. Characterization of additional *TIR1* polymorphisms.** Nonsynonymous polymorphisms in the Fbox domain of *TIR1* and with dN/dS value <1 were synthesized and co-expressed in yeast with fluorescently labeled IAA1 or IAA17. Degradation was assessed using flow cytometry on cultures exposed to different concentrations of the auxin, indole-3-acetic acid (IAA) for one hour. Error bars represent 95% confidence intervals around the median fluorescence calculated from three independent experiments. In many cases, intervals are small enough that they appear as a single line.

**S7 Fig. Characterization of additional AFB2 polymorphisms.** Nonsynonymous polymorphisms in the F-box domain of AFB2 and or with dN/dS values <1 were synthesized and co-expressed in yeast with fluorescently labeled IAA1 or IAA17. Degradation was assessed using flow cytometry on cultures exposed to different concentrations of the auxin, indole-3-acetic acid (IAA) for one hour. Error bars represent 95% confidence intervals around the median fluorescence calculated from three independent experiments. In many cases, intervals are small enough that they appear as a single line.

**S8 Fig. TIR1^Col^ accumulates to similar levels to polymorphic TIR1 variants in yeast.** Three replicate lysates of IAA17 and FLAG-tagged TIR1-variant expressing yeast strains were subjected to western blotting. A nonspecific band (*) is included as a loading control.

**S9 Fig. TIR1 variant expression in transgenic lines.** Triplicate RNA samples were isolated from T3 plants from each independent transformant line and subjected to reverse transcription and quantitative PCR for *TIR1* and *PP2A* as a control. Mean *TIR1* expression relative to *PP2A* normalized to the average expression across all *TIR1^Col^* and *TIR1^T154S^* lines is shown. Error bars represent SEM.

**S10 Fig. *tir1-10* is an amorph in synthetic auxin-induced degradation assays.** A yeast expression cassette recapitulating the *tir1-10* allele (TIR1^K159*^) was co-expressed with YFP-IAA17 as a homozygous diploid and along with full-length TIR1^Col^ and an empty expression cassette (null). Each yeast strain was treated with various concentrations of auxin for one hour during log-phase growth. YFP-IAA17 fluorescence was measured by flow cytometry. Mean fluorescence +/- SE calculated from four experiments are represented by points and error bars respectively. Some error bars are within the points.

**S11 Appendix. Supplemental information.** Complete analytical methods, detailed protocols and additional figures for each section of the main text.

## REFERENCES

Bella J., Hindle K. L., McEwan P. A., Lovell S. C., 2008 The leucine-rich repeat structure. Cell. Mol. Life Sci. 65: 2307–2333.

Calderón Villalobos L. I. A., Lee S., De Oliveira C., Ivetac A., Brandt W., et al., 2012 A combinatorial TIR1/AFB-Aux/IAA co-receptor system for differential sensing of auxin. Nat Chem Biol 8: 477–85.

Čermák T., Curtin S. J., Gil-Humanes J., Čegan R., Kono T. J. Y., et al., 2017 A Multipurpose Toolkit to Enable Advanced Genome Engineering in Plants. Plant Cell Online 29: 1196–1217.

Charif D., Lobry J. R., 2007 SeqinR 1.0-2: a contributed package to the R project for statistical computing devoted to biological sequences retrieval and analysis. In: Bastolla U, Porto M, Roman HE, Vendruscolo M (Eds.), Structural approaches to sequence evolution: Molecules, networks, populations, Biological and Medical Physics, Biomedical Engineering. Springer Verlag, New York, pp. 207–232.

Delker C., Pöschl Y., Raschke A., Ullrich K., Ettingshausen S., et al., 2010 Natural Variation of Transcriptional Auxin Response Networks in Arabidopsis thaliana. Plant Cell 22: 2184–2200.

Dezfulian M. H., Jalili E., Roberto D. K. A., Moss B. L., Khoo K., et al., 2016 Oligomerization of SCF TIR1 Is Essential for Aux/IAA Degradation and Auxin Signaling in Arabidopsis. PLOS Genet 12: e1006301.

Dharmasiri N., Dharmasiri S., Weijers D., Lechner E., Yamada M., et al., 2005a Plant Development Is Regulated by a Family of Auxin Receptor F Box Proteins. Dev. Cell 9: 109–119.

Dharmasiri N., Dharmasiri S., Estelle M., 2005b The F-box protein TIR1 is an auxin receptor. Nature 435: 441–445.

Dreher K. A., Brown J., Saw R. E., Callis J., 2006 The Arabidopsis Aux/IAA Protein Family Has Diversified in Degradation and Auxin Responsiveness. Plant Cell 18: 699–714.

Enders T. A., Strader L. C., 2015 Auxin activity: Past, present, and future. Am. J. Bot. 102: 180–196.

Engler C., Gruetzner R., Kandzia R., Marillonnet S., 2009 Golden Gate Shuffling: A One-Pot DNA Shuffling Method Based on Type IIs Restriction Enzymes. PLoS ONE 4: e5553.

Galli M., Liu Q., Moss B. L., Malcomber S., Li W., et al., 2015 Auxin signaling modules regulate maize inflorescence architecture. Proc. Natl. Acad. Sci.: 201516473.

Gibson D. G., Young L., Chuang R.-Y., Venter J. C., Hutchison C. A., et al., 2009 Enzymatic assembly of DNA molecules up to several hundred kilobases. Nat. Methods 6: 343–345.

Gray W. M., Pozo J. C. del, Walker L., Hobbie L., Risseeuw E., et al., 1999 Identification of an SCF ubiquitin-ligase complex required for auxin response in Arabidopsis thaliana. Genes Dev 13: 1678–91.

Grefen C., Donald N., Hashimoto K., Kudla J., Schumacher K., et al., 2010 A ubiquitin-10 promoter-based vector set for fluorescent protein tagging facilitates temporal stability and native protein distribution in transient and stable expression studies. Plant J. 64: 355–365.

Guilfoyle T. J., Hagen G., 2007 Auxin response factors. Curr. Opin. Plant Biol. 10: 453–460.

Guseman J. M., Hellmuth A., Lanctot A., Feldman T. P., Moss B. L., et al., 2015 Auxin-induced degradation dynamics set the pace for lateral root development. Development 142: 905–909.

Havens K. A., Guseman J. M., Jang S. S., Pierre-Jerome E., Bolten N., et al., 2012 A synthetic approach reveals extensive tunability of auxin signaling. Plant Physiol 160: 135–42.

Hellens R. P., Edwards E. A., Leyland N. R., Bean S., Mullineaux P. M., 2000 pGreen: a versatile and flexible binary Ti vector for Agrobacterium-mediated plant transformation. Plant Mol. Biol. 42: 819–832.

Hillson N. J., Rosengarten R. D., Keasling J. D., 2012 j5 DNA Assembly Design Automation Software. ACS Synth. Biol. 1: 14–21.

Hu Z., Keçeli M. A., Piisilä M., Li J., Survila M., et al., 2012 F-box protein AFB4 plays a crucial role in plant growth, development and innate immunity. Cell Res. 22: 777–781.

Lavy M., Estelle M., 2016 Mechanisms of auxin signaling. Development 143: 3226–3229.

Mathan J., Bhattacharya J., Ranjan A., 2016 Enhancing crop yield by optimizing plant developmental features. Development 143: 3283–3294.

Meijón M., Satbhai S. B., Tsuchimatsu T., Busch W., 2014 Genome-wide association study using cellular traits identifies a new regulator of root development in Arabidopsis. Nat. Genet. 46: 77–81.

Moss B. L., Mao H., Guseman J. M., Hinds T. R., Hellmuth A., et al., 2015 Rate Motifs Tune Auxin/Indole-3-Acetic Acid Degradation Dynamics. Plant Physiol. 169: 803–813.

Nei M., Gojobori T., 1986 Simple methods for estimating the numbers of synonymous and nonsynonymous nucleotide substitutions. Mol. Biol. Evol. 3: 418–426.

Nishimura K., Fukagawa T., Takisawa H., Kakimoto T., Kanemaki M., 2009 An auxin-based degron system for the rapid depletion of proteins in nonplant cells. Nat. Methods 6: 917–922.

Obenchain V., Lawrence M., Carey V., Gogarten S., Shannon P., et al., 2014 VariantAnnotation: a Bioconductor package for exploration and annotation of genetic variants. Bioinformatics 30: 2076–2078.

Parry G., Calderon-Villalobos L. I., Prigge M., Peret B., Dharmasiri S., et al., 2009 Complex regulation of the TIR1/AFB family of auxin receptors. Proc. Natl. Acad. Sci. 106: 22540–22545.

Pfeifer B., Wittelsbürger U., Ramos-Onsins S. E., Lercher M. J., 2014 PopGenome: An Efficient Swiss Army Knife for Population Genomic Analyses in R. Mol. Biol. Evol. 31: 1929–1936.

Pierre-Jerome E., Jang S. S., Havens K. A., Nemhauser J. L., Klavins E., 2014 Recapitulation of the forward nuclear auxin response pathway in yeast. Proc. Natl. Acad. Sci. 111: 9407–9412.

Pierre-Jerome E., Moss B. L., Lanctot A., Hageman A., Nemhauser J. L., 2016 Functional analysis of molecular interactions in synthetic auxin response circuits. Proc. Natl. Acad. Sci. U. S. A. 113: 11354–11359.

Pierre-Jerome E., Wright R. C., Nemhauser J., 2017 Characterizing Auxin Response Circuits in Saccharomyces cerevisiae by Flow Cytometry. In: Kleine-Vehn J, Sauer M (Eds.), *Plant Hormones*, Methods in Molecular Biology. Springer New York, pp. 271–281.

Prigge M. J., Greenham K., Zhang Y., Santner A., Castillejo C., et al., 2016 The Arabidopsis Auxin Receptor F-Box Proteins AFB4 and AFB5 Are Required for Response to the Synthetic Auxin Picloram. G3 GenesGenomesGenetics 6: 1383–1390.

Rasband W., 1997 *ImageJ*. U. S. National Institutes of Health, Bethesda, Maryland, USA.

Rosas U., Cibrian-Jaramillo A., Ristova D., Banta J. A., Gifford M. L., et al., 2013 Integration of responses within and across Arabidopsis natural accessions uncovers loci controlling root systems architecture. Proc. Natl. Acad. Sci. 110: 15133–15138.

Ruegger M., Dewey E., Gray W. M., Hobbie L., Turner J., et al., 1998 The TIR1 protein of Arabidopsis functions in auxin response and is related to human SKP2 and yeast Grr1p. Genes Dev. 12: 198–207.

Sambrook J., Russell D. W., 2001 *Molecular Cloning: A Laboratory Manual*. CSHL Press.

Schmitz R. J., Schultz M. D., Urich M. A., Nery J. R., Pelizzola M., et al., 2013 Patterns of population epigenomic diversity. Nature 495: 193–198.

Sun C., Wang B., Wang X., Hu K., Li K., et al., 2016 Genome-Wide Association Study Dissecting the Genetic Architecture Underlying the Branch Angle Trait in Rapeseed (Brassica napus L.). Sci. Rep. 6: 33673.

Tan X., Calderon-Villalobos L. I. A., Sharon M., Zheng C., Robinson C. V., et al., 2007 Mechanism of auxin perception by the TIR1 ubiquitin ligase. Nature 446: 640–5.

Terrile M. C., París R., Calderón-Villalobos L. I. A., Iglesias M. J., Lamattina L., et al., 2012 Nitric oxide influences auxin signaling through S-nitrosylation of the Arabidopsis TRANSPORT INHIBITOR RESPONSE 1 auxin receptor. Plant J. Cell Mol. Biol. 70: 492–500.

Walsh T. A., Neal R., Merlo A. O., Honma M., Hicks G. R., et al., 2006 Mutations in an Auxin Receptor Homolog AFB5 and in SGT1b Confer Resistance to Synthetic Picolinate Auxins and Not to 2,4-Dichlorophenoxyacetic Acid or Indole-3-Acetic Acid in Arabidopsis. Plant Physiol. 142: 542–552.

Yu H., Moss B. L., Jang S. S., Prigge M., Klavins E., et al., 2013 Mutations in the TIR1 Auxin Receptor That Increase Affinity for Auxin/Indole-3-Acetic Acid Proteins Result in Auxin Hypersensitivity. Plant Physiol. 162: 295–303.

Yu H., Zhang Y., Moss B. L., Bargmann B. O. R., Wang R., et al., 2015 Untethering the TIR1 auxin receptor from the SCF complex increases its stability and inhibits auxin response. Nat. Plants 1: 14030.

Zhang X., Henriques R., Lin S., Niu Q., Chua N., 2006 Agrobacterium-mediated transformation of Arabidopsis thaliana using the floral dip method. Nat. Protoc. 1: 641–6.

Zhang L., Ward J. D., Cheng Z., Dernburg A. F., 2015 The auxin-inducible degradation (AID) system enables versatile conditional protein depletion in C. elegans. Dev. Camb. Engl.

